## Supplement for "A benchmarked comparison of software packages for time-lapse image processing of monolayer bacterial population dynamics"

### Supporting Information Text

#### 1. Image processing package overviews

**Table S1. Features of the software packages evaluated in this study**

| Feature | CellProfiler | SuperSegger | DeLTA | FAST |
|---|---|---|---|---|
| Pre-processed inputs | ✓ | × | × | ✓ |
| GUI | ✓ | ✓ | × | ✓ |
| Online segmentation correction | ✓ | × | × | × |
| Online tracking correction | × | × | × | ✓ |
| Cell type | eukaryote/prokaryote | prokaryote | eukaryote/prokaryote | prokaryote |
| Language | python | MATLAB/C++ | python | MATLAB |
| Release date | 2006 | 2016 | 2020 | 2021 |
| Online discussion forum | ✓ | × | ✓ | × |

#### 2. Results

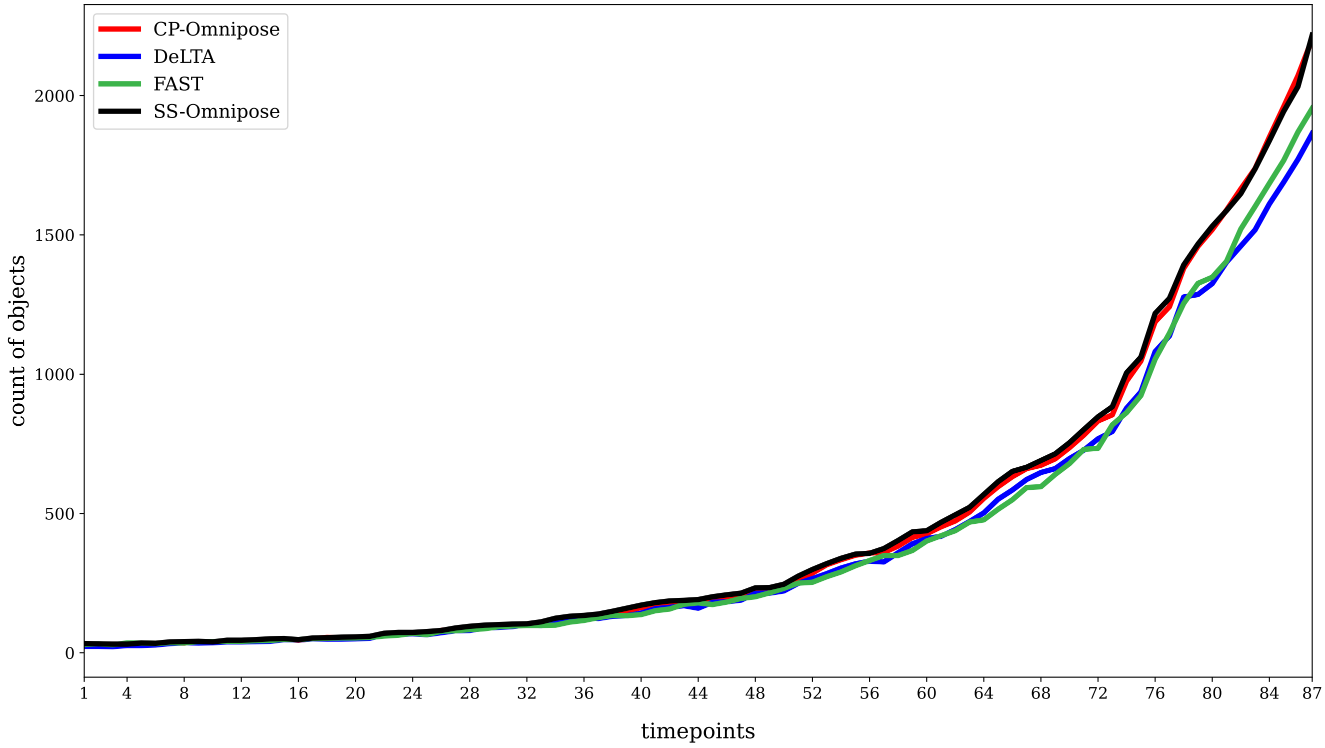


**Fig. S1.** Segmentation object count results for time-lapse of unconstrained *X. campestris* growth. Ground truth was not generated for this dataset. The time interval between each frame is 3 minutes, and the total duration of the experiment is 258 minutes. Full dataset at Github (CP: CellProfiler, SS: SuperSegger)

##### A. Segmentation object counts for *X. campestris*.

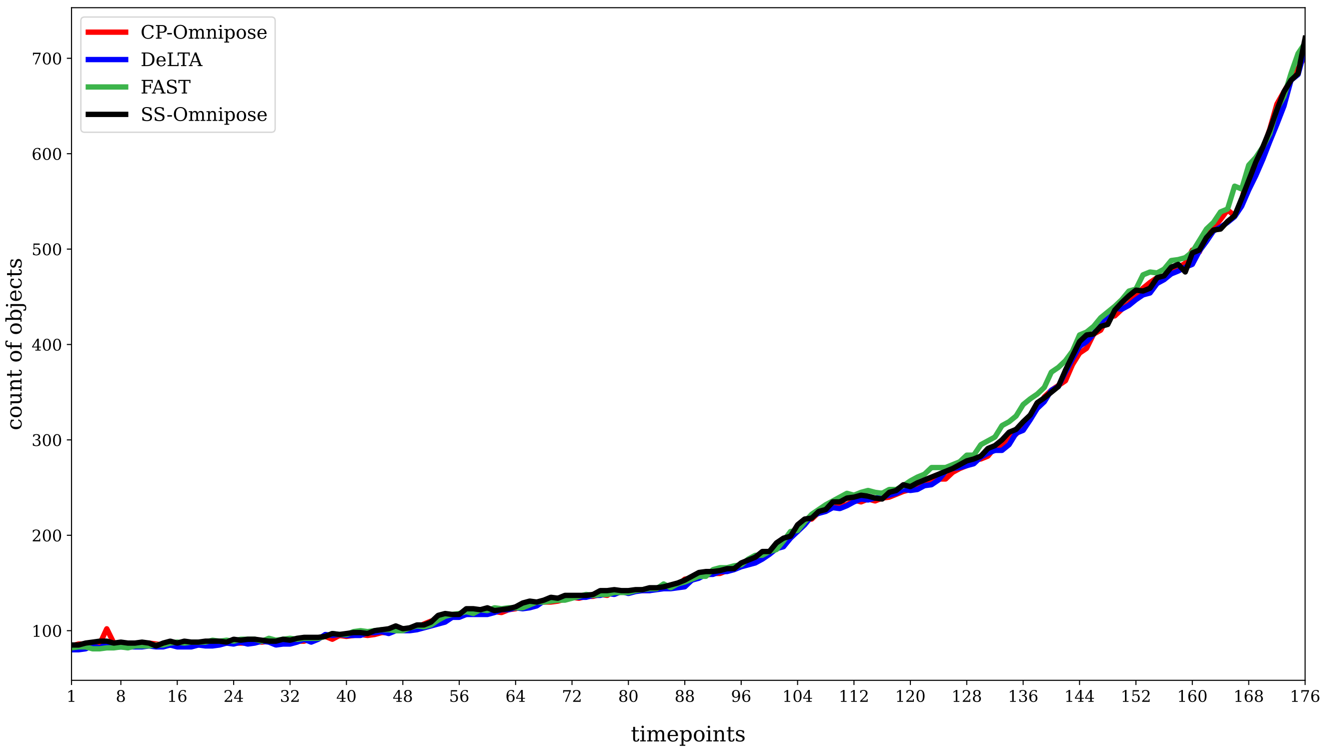


**Fig. S2.** Segmentation object count results for time-lapse of unconstrained *P. putida* growth. Ground truth was not generated for this dataset. The time interval between each frame is 3 minutes, and the total duration of the experiment is 528 minutes. Full dataset at Github. (CP: CellProfiler, SS: SuperSegger)

#### B. Segmentation object counts for *P.putida*.

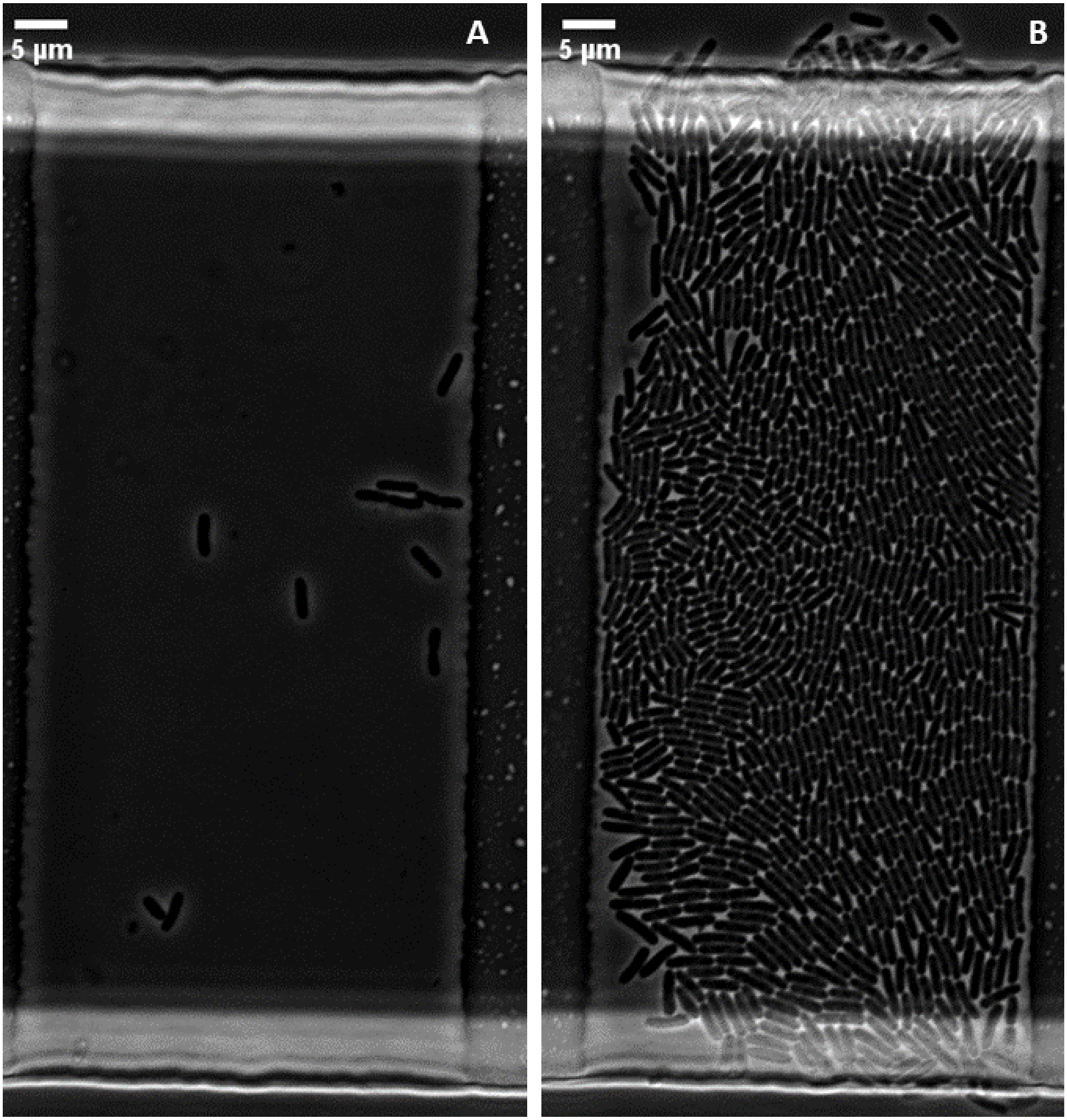


**Fig. S3.** *E.* coli growth in a constrained environment. (A) First and (B) last (54th) timestep of *E.coli* growth in a microfluidic trap. The initial frame contains 10 cells. The population grows to 1049 cells by the 54th timestep, after 162 minutes. A distinct halo of increased brightness can be observed near the chamber entrances; this can be attributed to imaging artifacts arising from the neighbouring (taller) flow channel.

#### C. First and last timepoints of microfluidic time-lapse.

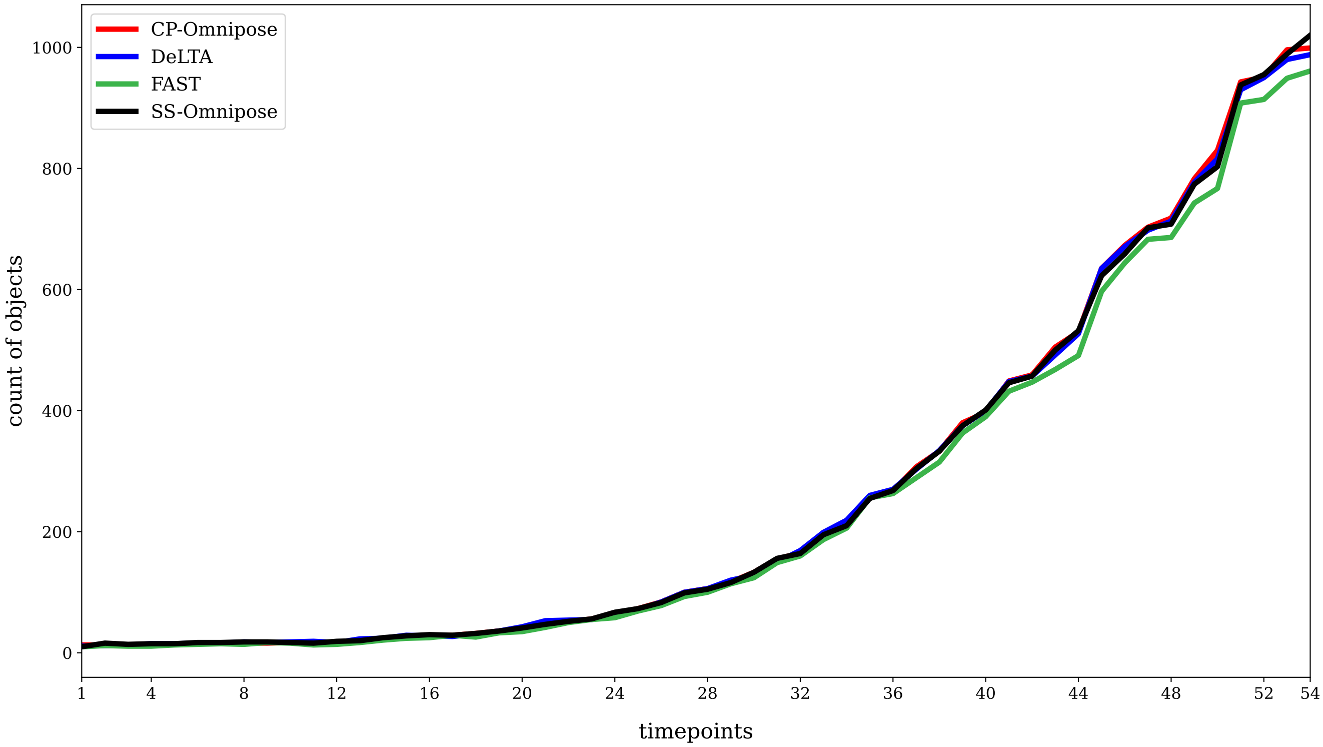


**Fig. S4.** Segmentation object count results for time-lapse of constrained *E.coli* growth. Ground truth was not generated for this dataset. The time interval between each frame is 1.5 minutes, and the total duration of the experiment is 81 minutes. (CP: CellProfiler, SS: SuperSegger)

9 ## D. Segmentation object counts for *E.coli* in the microfluidic trap.

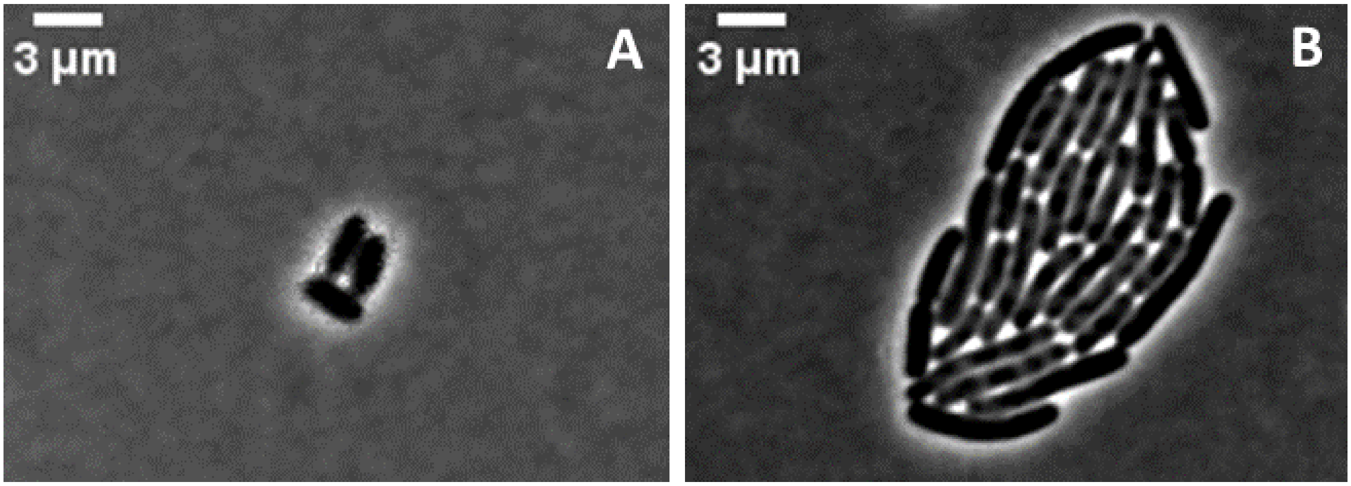


**Fig. S5.** (A) First and (B) last (31st) frame of a cropped time-lapse of unconstrained *E.coli* growth. The time interval between each frame is 5 minutes, and the total duration of the experiment is 150 minutes.

10 ## E. Representative images from the workable *E.coli* time-lapses.

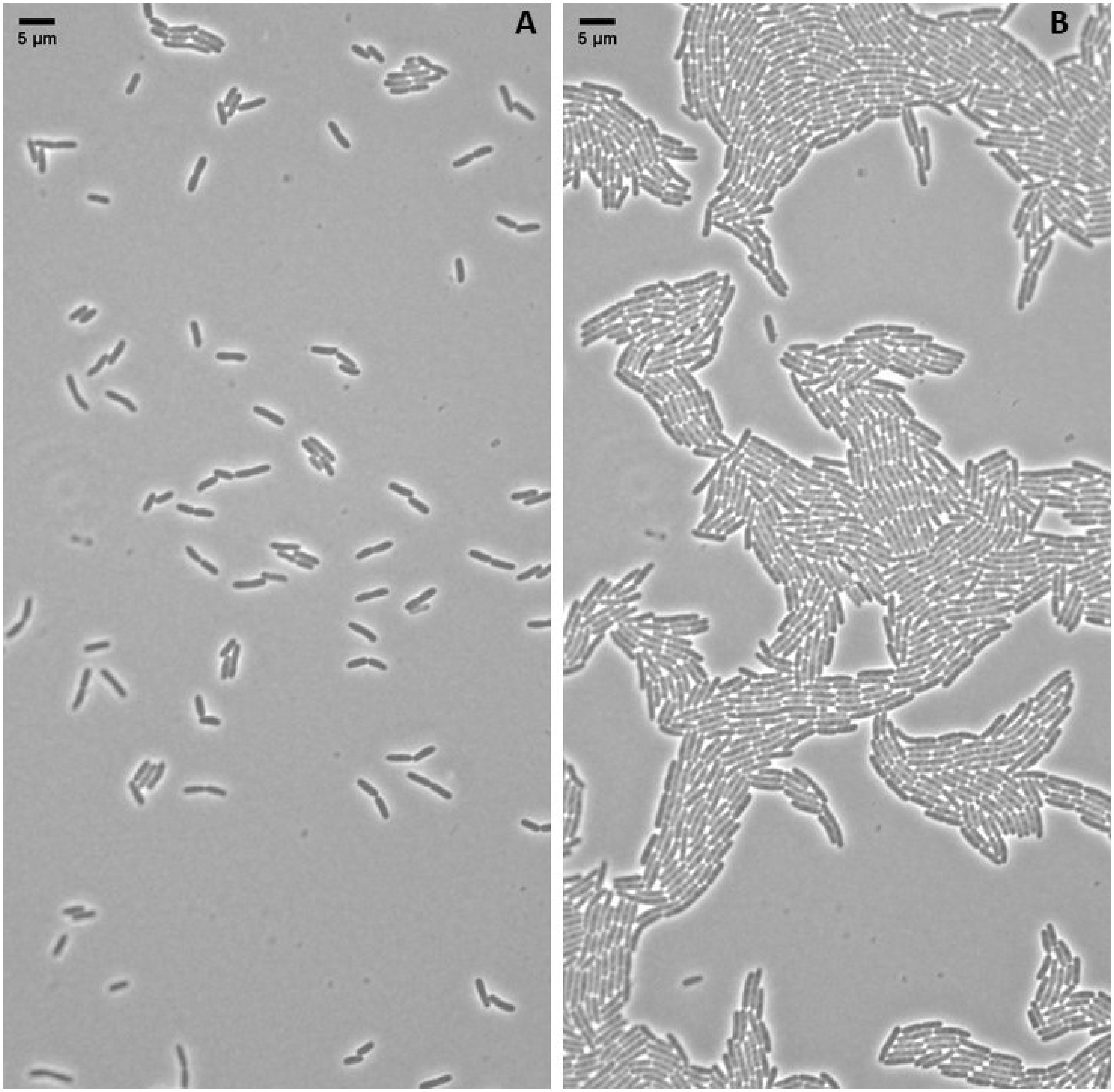


**Fig. S6.** (A) First and (B) last (100th) images in a time-lapse of unfiltered *E.coli* growth in an unconstrained environment. The initial frame contains 105 cells. The population grows to 1140 by the final frame. Cells touching the edges are not counted. The cells are in monolayer throughout 150 minute duration

#### F. Unfiltered *E.coli* time-lapses.
